# Live cell single molecule tracking and localization microscopy of bioorthogonally labeled plasma membrane proteins

**DOI:** 10.1101/660118

**Authors:** Andres I. König, Raya Sorkin, Ariel Alon, Dikla Nachmias, Kalyan Dhara, Guy Brand, Ofer Yifrach, Eyal Arbely, Yael Roichman, Natalie Elia

## Abstract

Tracking the localization and mobility of individual proteins in live cells is key for understanding how they mediate their function. Such information can be obtained from single molecule imaging techniques such as Single Particle Tracking (SPT) and Single Molecule Localization Microscopy (SMLM). Genetic code expansion (GCE) combined with bioorthogonal chemistry offers an elegant approach for direct labeling of proteins with fluorescent dyes, holding great potential for improving protein labeling in single molecule applications. Here we calibrated conditions for performing SPT and live-SMLM of bioorthogonally labeled plasma membrane proteins in live mammalian cells. Using SPT, the diffusion of bioorthogonally labeled EGF receptor and the prototypical Shaker voltage-activated potassium channel (Kv) was measured and characterized. Applying live-SMLM to bioorthogonally labeled Shaker Kv channels enabled visualizing the plasma membrane distribution of the channel over time with ~30 nm accuracy. Finally, by competitive labeling with two Fl-dyes, SPT and live-SMLM were performed in a single cell and both the density and dynamics of the EGF receptor were measured at single molecule resolution in subregions of the cell. We conclude that GCE and bioorthogonal chemistry is a highly suitable, flexible approach for protein labeling in quantitative single molecule applications that outperforms current protein live-cell labeling approaches.

## INTRODUCTION

Genetic code expansion (GCE) together with biorthogonal labeling offers an elegant tool for labeling proteins with organic Fluorescent dyes (Fl-dyes) in live mammalian cells. In GCE-based labeling, a non-canonical amino acid (ncAA) carrying a functional group is incorporated into the protein sequence in response to an in-frame amber stop codon (TAG), via an orthogonal tRNA/tRNA-synthetase pair (reviewed in ^1,2^). Labeling is then carried out by a rapid and specific bioorthogonal reaction between the functional group and a Fl-dye. As a result, the protein of interest is fluorescently labeled on a specific residue, avoiding the need for conjugating protein tag fusions (i.e. fluorescent proteins or self-labeling proteins) in live cell applications.

Direct labeling of proteins in live cells with Fl-dyes has several advantages for quantitative, high-end live cell imaging. First, Fl-dyes are bright and photostable and therefore cells can be imaged for longer times and at reduced laser intensities ^3^. Second, the label itself is relatively small (Fl-dye, ~0.5 nm; GFP, 4.2 nm; antibodies >10nm; quantum dots 2-60 nm) thereby allowing higher accuracy in localization measurements and better representation of the physiological properties of the protein ^3–5^. Third, labeling does not involve amplification, and therefore the number of molecules can be quantified. Fourth, the Fl-dye is applied at the last step of the reaction providing flexibility in optimizing the Fl-dye to specific applications and offering simultaneous labeling of protein populations with more than one dye ^6^. Indeed, GCE and bioorthogonal labeling have been demonstrated, in recent years, in several microscopy techniques including live cell imaging and the super resolution techniques Structured Illumination Microscopy (SIM), Stimulated Emission Depletion (STED) and Single Molecule Localization Microscopy (SMLM) ^6–11^. Yet, these studies focused on the proof of concept and very little quantitative information on the spatiotemporal dynamics of proteins in cells has been obtained from them.

Here we set out to perform quantitative analysis of the dynamics and spatial distribution of plasma membrane (PM) proteins labeled via GCE and bioorthogonal chemistry. To this aim, we calibrated conditions for performing the live-cell single-molecule applications, Single Particle Tracking (SPT) and live-SMLM using extracellular bioorthogonal labeling. We then applied our optimized SPT assay, to measure the diffusion coefficients of bioorthogonally labeled epidermal growth factor receptor (EGFR) and the prototypical Shaker B voltage dependent potassium (Kv) channel. Significantly more tracks and longer trajectories were obtained for bioorthogonally labeled epidermal growth factor receptor (EGFR), compared to EGFR-GFP. Moreover, while SPT was successfully applied to bioorthogonally-labeled Shaker B, no tracks could be generated using mCherry-Shaker B. Consistent with previous reports, EGFR diffusion was more confined upon ligand activation and Shaker B diffusion became less confined in the presence of the actin polymerization inhibitor Latrunculin A (LatA) ^12–15^. Together these results indicate that labeling proteins via GCE and bioorthogonal chemistry outperforms Fl-protein labeling for particle tracking. SMLM experiments of bioorthogonally labeled EGFR and Shaker B were successfully performed in fixed and live cells using our calibrated assay and allowed quantifying the dynamic distribution of PM proteins over time. Finally, using simultaneous labeling with two different Fl-dyes we performed live-SMLM and SPT in a single cell and determined the density and the dynamics of EGFR in subcellular regions at nanoscale resolution. Based on these results we concluded that GCE-based bioorthogonal labeling of proteins is an improved, more flexible approach for quantitative live cell single molecule applications.

## RESULTS

### Calibrating conditions for SPT of bioorthogonally labeled PM proteins using EGFR

We began by testing the applicability of bioorthogonal labeling via GCE for single molecule imaging by using EGFR, a well-studied receptor that has been previously labeled via GCE, as a benchmark ^16^. To this end, EGFR was mutated to carry a TAG codon at the previously published labeling site, Leu 128, and cloned into a single expression vector that encodes the cognate pair of tRNA_cua_:tRNA-synthetase ^17,18^ (Fig. S1, top panel). The vector was then expressed in COS7 cells in the presence of the ncAA Bicyclo[6.1.0]nonyne-L-lysine (BCNK), which reacts bioorthogonally with tetrazine conjugated dyes (Fig. 1a, left panel) ^16,19^. Under these conditions, specific PM labeling was obtained using the cell impermeable tetrazine conjugated Fl-dyes: AF647, Cy3, Cy5 or ATTO532, with Tet-Cy3 providing the most robust labeling, indicating that the EGFR^128BCNK^ is efficiently expressed and targeted to the PM (Fig. 1a). Next, we tested the suitability of the different dyes for SPT, in vitro. This analysis was required because the tetrazine moiety is known to change the fluorogenicity of Fl-dyes and therefore may also affect their photophysical properties ^2^. Among the Tet-conjugated dyes tested, Tet-Cy3 had the best overall properties for SPT applications; it was the brightest and it produced the longest tracks and the largest number of tracks (Fig. S1b).

**Fig. 1.**
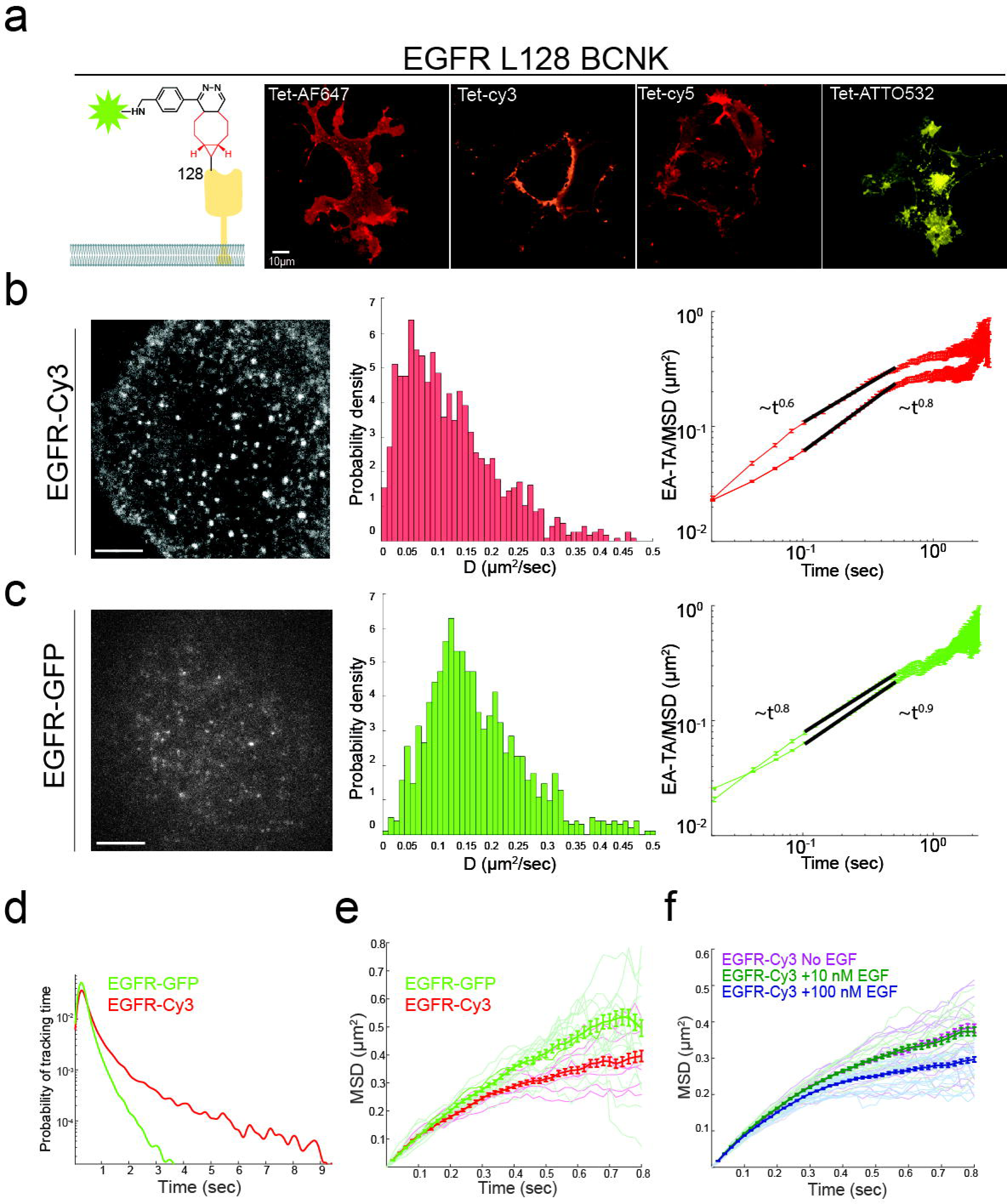
Bioorthogonal labeling with Fl-dyes enables single particle tracking of EGFR. **(a)** Left panel: schematic representation of the labeling strategy. The extracellular domain of EGFR (yellow) was mutated to carry the ncAA BCNK (red) at position 128. Labeling was obtained via a bioorthogonal reaction between BCNK and a tetrazine conjugated Fl-dye (green star). Panels 2-5: Live COS7 cells expressing EGFR^128BCNK^ together with GCE components (Fig. S1a) were labeled with 1.5 µM of: Tet-AF647, Tet-Cy3, Tet-Cy5 or Tet-ATTO532 and imaged using a spinning disk confocal microscope. Shown are single slices taken from the center of the cell. Scale bar = 10 µm. **(b, c)** COS7 cells transfected with either GCE system plasmid with EGFR^128BCNK^ and labeled with Tet-Cy3 or EGFR-GFP, were imaged in TIRF mode at 50 fps. Tracks were then obtained as described in methods section. Left panel: representative images of the first frame (the complete video sequences are provided in Videos 1 and 2). Middle panels: The diffusion coefficient [D] of all particles tracked. EGFR-Cy3 median, 0.11 µm^2^/s (with 95% confidence interval of 0.02-0.4 µm^2^/s, n=1177). EGFR-GFP median, 0.15 µm^2^/s (with 95% confidence interval of 0.04-0.47 µm^2^/s, n=1018). Right panels: ensemble-averaged MSD (top plots) and ensemble-averaged time-averaged MSD (bottom plots) in log scale, MSD∼t^α^, where α is the anomalous exponent **(d)** The tracking time probability density function of EGFR-GFP (green) or EGFR-Cy3 (red). **(e)** MSD curves of EGFR-GFP (green) or EGFR-Cy3 (red) as a function of time. Thick lines are the ensemble means for each labeling type with error bars representing SEM. Thin lines represent measurements obtained from individual cells (EGFR-Cy3 n=9 cells, EGFR-GFP n=17 cells, obtained from three independent experiments). **(f)** MSD curves obtained for EGFR-Cy3 under three different concentrations of EGF: no EGF (magenta), 10 nM EGF (dark green) or 100 nM EGF (blue) as a function of time. Thick lines are the ensemble means for each labeling type with error bars representing SEM. Thin lines represent measurements obtained from individual cells (no EGF n=18 cells, 10 nM EGF n=15 cells, and 100 nM EGF n=18 cells).

A uniform decoration of the PM was observed in cells expressing EGFR^128BCNK^ labeled with Tet-Cy3 and imaged in total internal reflection (TIRF) mode, which supports high labeling efficiencies, but does not allow single molecules imaging. Under mild photobleaching conditions, single spots could be readily detected with relatively high signal to noise ratio (SNR) (2.66 ± 0.45) (Fig. 1b). In cells expressing EGFR-GFP individual spots could be seen without applying photobleaching and were significantly dimmer (SNR = 2.01 ± 0.27), indicating that bioorthogonally labeled EGFR is brighter and more efficiently expressed at the PM compared to EGFR-GFP (Fig. 1b, c and videos 1 and 2).

Next, we plotted the distribution of brightness measured for individual foci in single frames and fitted it to the smallest sum of Gaussian distributions that fit our data well (Fig. S2). For EGFR-Cy3 the best fit was obtained for three Gaussians, suggesting that one to three labeled EGFR molecules can reside in a single diffraction limited spot. For EGFR-GFP, on the other hand, the best fit was obtained for a single Gaussian, indicating a homogenous population. EGFR is known to reside in monomers, dimers and small clusters at the PM ^14^. It is therefore reasonable that at high expression levels labeled EGFR will mix with the endogenous population of EGFR at different stoichiometries giving rise to heterogeneous brightness population, as observed for EGFR-Cy3. The results obtained for EGFR-GFP may suggest that EGFR-GFP only resides in monomers at the PM or that under these expression levels, on average, one or less EGFR-GFP molecules assemble in each EGFR complex.

For tracking analysis, trajectories were extracted from videos of EGFR tagged with GFP (EGFR-GFP) or bioorthogonally labeled with Cy3 (EGFR-Cy3), recorded in TIRF mode at 20 ms intervals for up to three minutes (Fig. 1b, c). For each condition, videos recorded in at least three different independent experiments were used for the analysis. About 810 trajectories were obtained from each cell labeled via GCE (EGFR-Cy3) while only ~290 trajectories were obtained from cells expressing EGFR-GFP. This difference may result from the tendency of Fl-dyes to nonspecifically bind to membranes and to the glass bottom of the imaging chamber ^3,20^. To avoid including these nonspecific stationary Fl-dye molecules in our measurements, we tracked fluorophores located outside the cells and calculated their mean square displacements (MSDs) and diffusion rates (Fig. S3a, c). We then used these background trajectories to filter out trajectories with similar MSD values (Fig. S3e). To avoid bias, a similar procedure was applied to EGFR-GFP tracks (Fig. S3b, d, f). Additionally, we noticed that tracks obtained for bioorthogonally labeled EGFR were considerably longer than EGFR-GFP tracks, allowing measuring diffusion at longer timescales (Fig. 1d). After filtering background trajectories and short trajectories (<20 frames) about 130 trajectories were obtained from each cell with EGFR labeled via GCE (EGFR-Cy3) and only ~60 trajectories were obtained from cells expressing EGFR-GFP. The larger number of tracks with increased length obtained for bioorthogonally labeled EGFR improves the accuracy of the measurements. Therefore, measurements obtained using EGFR-Cy3 are likely to provide more realistic view of EGFR dynamics in cells.

Under these conditions, MSDs and median diffusion rates were calculated for EGFR-GFP (0.15 µm^2^/s with 95% confidence interval of 0.04-0.47 µm^2^/s) and for bioorthogonally labeled EGFR (0.11 µm^2^/s with 95% confidence interval of 0.02-0.4 µm^2^/s) (Fig, 1b, c, e). Fluorescence recovery after bleaching (FRAP) analysis performed in cells expressing EGFR^128BCNK^ and bioorthogonally labeled with Cy3 resulted in a mean diffusion coefficient of 0.05 µm^2^/s +-0.017, which falls in the range of diffusion rates measured by SPT (Fig. S4a). To further characterize EGFR diffusion, we plotted the ensemble-averaged and time-averaged MSDs ^21^. The observed scaling exponents corresponded to subdiffusion, e.g. with anomalous diffusion exponents α<1 ^22^. Different curves were obtained for ensemble and time averaged MSDs of bioorthogonally labeled EGFR, supporting non-ergodic behavior at short time lags (Fig. 1b, right panel). A milder difference was observed for EGFR-GFP (Fig. 1c, right panel). The time-averaged MSD (TA-MSD) plots as a function of lag time Δ for individual trajectories are shown in figure S4b. Plotting the turning angle distribution showed a high probability around turning angle of 180 degrees (π radians), which points to strong directional correlation in the particle motion for EGFR-Cy3; specifically, particles tend to reverse their direction after each step. This is consistent with either motion in viscoelastic media or in confined geometries ^23^. The non-ergodicity at short timescales deduced from the MSD curves is more consistent with confined motion ^21,22,24–26^. Based on the ensemble averaged MSD plots, the time-averaged MSD plots and the turning probability plots, diffusion appeared to be noisier and less confined for EGFR-GFP (Fig. 1e, S4b, c).

EGFR is known to dimerize, cluster and endocytosed upon activation with its ligand, EGF ^14^. To quantify the changes in EGFR diffusion upon ligand activation we performed SPT of bioorthogonally labeled EGFR (EGFR-Cy3) in cells under starvation (no EGF) and in cells that were subjected to low (10 nM) and high (100 nM) doses of EGF. In agreement with previous reports, EGFR diffusion was slower and more confined upon EGFR activation (high doses of EGF) (Fig. 1f, right panel) ^12,14^.

### Measuring PM diffusion of the prototypic Kv channel, Shaker B, using bioorthogonal labeling and SPT

Convinced by the suitability of GCE-based labeling for SPT, we set out to study the dynamics of the prototypic Shaker B Kv channel. Voltage activating potassium channels are allosteric pore-forming proteins that open and close in response to changes in membrane potential ^27^. Kv channels are homotetrameric with the four channel subunits giving rise to the ion conductive pore domain ^28^. Each subunit is comprised of six transmembrane helices, with helices 1-4 (S1-S4) constructing the voltage-sensing domain, and helices 5 and 6 making the selective pore domain (Fig. 2a) ^27^. The channel is known to interact with scaffold proteins such as PSD-95 that determines the organization of the channel on the PM ^29,30^. Tagging the channel with fluorescent proteins is challenging because its N and C termini are respectively engaged in the fast inactivation and PM organization of the channel ^31,32^.

**Fig. 2.**
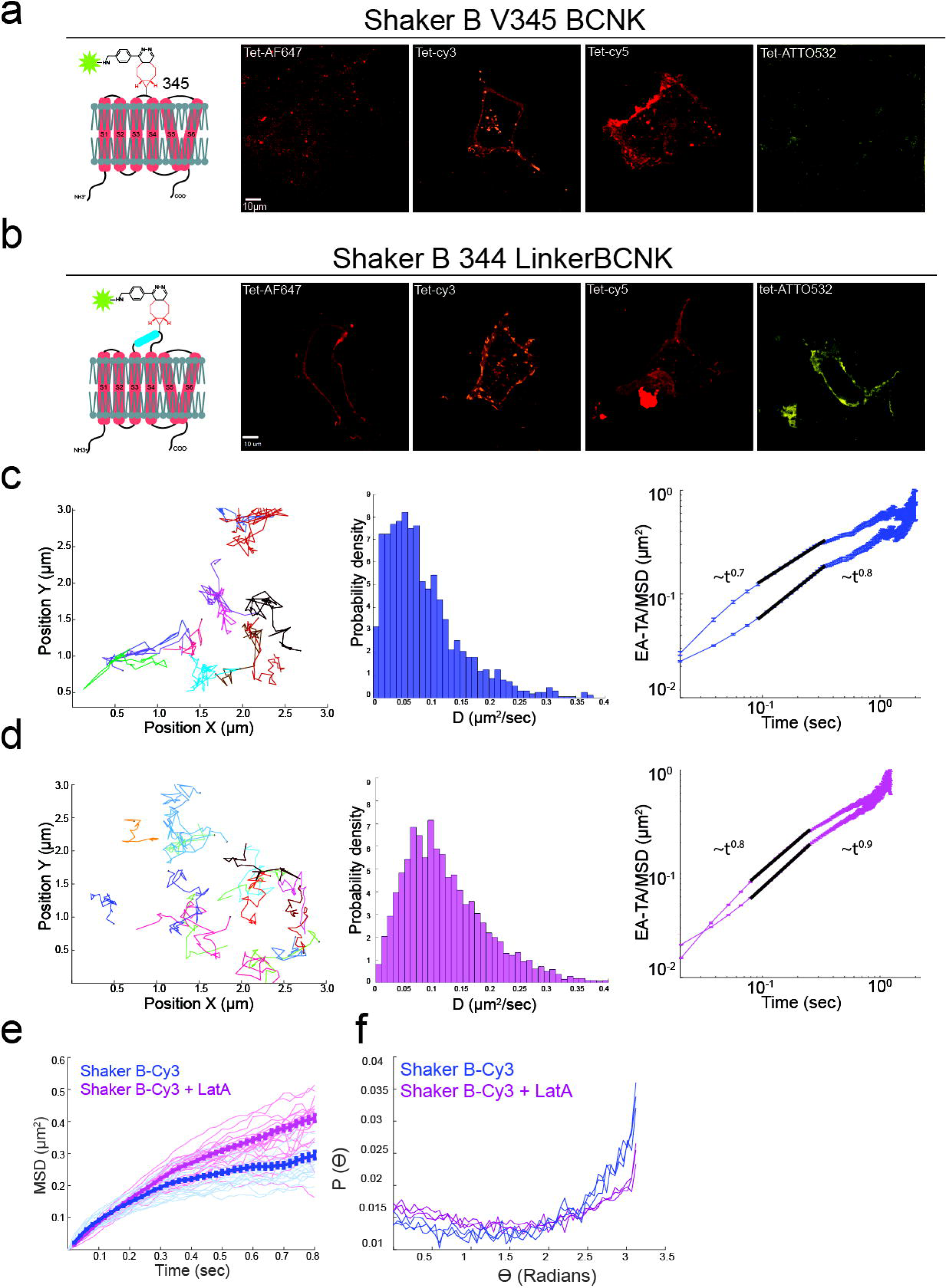
Measuring the PM dynamics of bioorthogonally labeled Shaker B channels. **(a, b)** COS7 cells transfected with the GCE system plasmid carrying Shaker B^V345BCNK^(**a**) or with Shaker B in which BCNK is encoded via a short 14 AA linker (**b**) (Fig. S1a), were labeled with the indicated tetrazine conjugated Fl-dye and imaged in a confocal spinning disk microscope. Shown are single slices taken from the center of the cell. Schematic representations of the labeling strategies are shown on the left. Scale bar = 10 µm. **(c, d)** Single particle tracking of Shaker B-Cy3 in naïve cells (**c**) and cells treated with 1 µM Latrunculin A (LatA) (**d**). Left panels, tracks obtained in representative COS7 cells. Each track is represented by a different color. Middle panels: the distribution of diffusion coefficients [D] of all particles tracked. Naïve cells median, 0.07 µm^2^/s (with 95% confidence interval of 0.01-0.3 µm^2^/s, n=1497). LatA treated cells median, 0.11 µm^2^/s (with 95% confidence interval of 0.02-0.4 µm^2^/s, n=4619). Right panels: ensemble averaged MSD (top plots) and ensemble averaged time-averaged MSD (bottom plots) in log scale, MSD∼t^α^, where α is the anomalous exponent (**e**) MSD curves of cells expressing either Shaker B-Cy3 (blue) or Shaker B-Cy3 with LatA (purple) as a function of time. Thick lines are the ensemble means for each labeling type with error bars representing SEM. Thin lines represent measurements obtained from individual cells (Shaker B-Cy3 n=16 cells, Shaker B-Cy3 with LatA n=27 obtained from three independent experiments). (**f**) Turning angle distributions for Shaker B-Cy3 (blue) and Shaker B-Cy3 + LatA (purple) measured with lag times of 1, 2 and 5 frames (20, 40 and 100 ms).

We began by testing the suitability of several extracellular positions in Shaker B for bioorthogonal labeling. Shaker B channels are not endogenously expressed in COS7 cells and therefore successful labeling of the channel on the PM will only be obtained upon tertramerization of four ncAA-containing Shaker B monomers ^27,28^. Two positions in the loop linking helices S1-S2, K253 and I275, and one in the S3-S4 loop, V345, were tested. While almost no labeling was obtained in cells expressing Shaker B^K253BCNK^ and Shaker B^I275BCNK^, robust and specific PM labeling was obtained by encoding BCNK at position 345 in Shaker B (Fig. 2a and S5). Interestingly, successful labeling of Shaker B^V345BCNK^ was obtained using Tet-Cy3 or Tet-Cy5, while no labeling obtained using Tet-ATTO-532 or Tet-AF647 (Fig. 2a). We suspected that this difference stems from the different chemical properties of the dyes and the proximity of the S3-S4 loop to the PM ^33,34^. To test this possibility, we inserted a 14-residue long peptide linker that encodes for BCNK, which we have recently optimized for biorthogonal labeling (unpublished results, Fig. S1a, bottom panel), between positions 344 and 345 in Shaker B. Using the linker, specific PM labeling of Shaker B was obtained using any of the dyes tested, suggesting that displacing BCNK from the PM may improve labeling (Fig. 2b). The linker-based version of BCNK-Shaker B (Shaker B^linkerBCNK^) was therefore used in all future experiments.

SPT experiments were successfully performed for Shaker B, bioorthogonally labeled with Tet-Cy3. The distribution of brightness best fitted to a single Gaussian (Fig. S6a). Because the channel is heterologously expressed and is known to be a homotetramer, we consider each fluorescence spot as a single channel labeled with four Cy3 molecules. Approximately 100 tracks were obtained from each cell, after filtering, with a median track length of 37 frames (compared to 130 tracks and median track length of 30 frames obtained for EGFR-Cy3), indicating that measurements are reliable. No trajectories could be generated upon applying similar experimental conditions to cells expressing mCherry-Shaker B due to low SNR (Figure S6b). This indicates that in addition to the potential artifacts associated with tagging the channel at the N terminal, the resulting signal is too dim for particle tracking. The median diffusion rate obtained for Shaker B-Cy3 after thresholding was 0.07 µm^2^/s (with a 95% confidence interval of 0.01-0.3 µm^2^/s) (Fig. 2c and S6c, d). This value is 0.04 µm^2^/s slower than the diffusion rate obtained for EGFR-Cy3, indicating that the channel is 36% less mobile than the receptor. Applying the actin polymerization inhibitor, LatA to cells with bioorthogonally labeled Shaker B, resulted in an increased diffusion rate (0.11 µm^2^/s with a 95% confidence interval of 0.02-0.4 µm^2^/s) (Fig. 2d). Additionally, based on the ensemble MSD plots and the turning angle probability plots, diffusion was less confined in the presence of LatA (Fig. 2d, e, f and S6e). This indicates that, in agreement with previous results, the actin cytoskeleton plays a role in controlling the mobility of the Shaker B channel on the PM ^15,35^.

### Performing live-SMLM on bioorthogonally labeled PM proteins

SMLM can be used to resolve the spatial distribution of PM proteins at ~30 nm resolution ^36–38^. Bioorthogonal labeling is expected to be advantageous for SMLM; the small size of the label improves the theoretical resolution of the technique and the superb photophysical properties of Fl-dyes enables acquisitions at reduced laser powers that may enable performing SMLM in live cell ^3,20^. High density labeling is required in SMLM and especially in live-SMLM experiments for obtaining a “true” representation of protein distributions. Encouraged by the high labeling densities obtained for EGFR and Shaker B in SPT experiments, we set to optimize conditions for live-SMLM of bioorthogonally labeled PM proteins. SMLM experiments were successfully performed in fixed COS7 cells expressing either EGFR^128BCNK^ or Shaker B^linkerBCNK^ and labeled with Tet-AF647 (Fig. S7).

More than 3,170 ± 553 localizations/µm^2^ and 1,600 ± 583 localizations/µm^2^ were obtained for EGFR and the Shaker Kv channel respectively, with an average localization uncertainty of 28.5 ±10.2 nm for EGFR and 23.3 ±6.9 nm for Shaker B. Approximately 195 localizations/µm^2^ and 318 localizations/µm^2^ were measured outside cells expressing EGFR-Cy3 and Shaker B, which corresponds to 6% and 20% background levels, respectively. The localization pattern of both EGFR and Shaker B on the PM appeared to be uniform with small accumulations appearing occasionally. To ensure that these accumulations are not an artifact resulting from low laser intensities we performed SMLM on fixed cells expressing EGFR^128BCNK^ and labeled with Tet-AF647 using 10 - 100% laser powers. We found that under low laser powers, EGFR was almost exclusively found in dense clusters, manifesting the previously reported artifacts ^39^. Increasing the laser power to 70% or above, abolished these artifacts indicating that measurements performed at these laser powers are reliable.

Next, we calibrated conditions for live SMLM. First, we tested different SMLM buffers that are known to induce efficient blinking of Tet-AF647 and are compatible with live cell imaging ^37,40^. We found OptiMEM supplemented with 5% glucose, a GLOXY oxygen scavenging system and 1% β-mercaptoethanol, suitable for extracellular live-SMLM. Second, we calibrated the acquisition protocol. In classical SMLM experiments, the number of new molecules that appear in each frame decreases overtime until the entire population of molecules is exhausted. Yet, to acquire representative data over time the number of localizations should be as similar as possible in each time point. By reducing the laser power to 70% and using a milder protocol for the 405-activation laser we obtained an acquisition protocol that is compatible with live cell recordings (Fig. S8). Last, we estimated temporal resolution. An SMLM image is built from a series of images hence, the temporal resolution should be set by the minimal number of frames needed for obtaining a representative image of the distribution of molecules across the PM. Based on SMLM experiments performed on bioorthogonaly-labeled Shaker Kv channels in fixed cells, we estimated that ~150 frames are needed for getting a representative SMLM image, providing a temporal resolution of 2.7 seconds (at 18 ms frame rate) (Fig. S9).

Using these optimized conditions, live-SMLM experiments were successfully recorded in COS7 cells expressing GCE modified Shaker B channels and labeled with Tet-AF647 (Fig. 3, video 3). Live SMLM videos of Shaker B-AF647 were recorded for 27 seconds and each 150 frames were merged into a single timepoint. Approximately 33 localizations/µm^2^ (total of 1577 localizations/cell) with an average uncertainty of 25.5±6.8 nm, were obtained in each timepoint. Under these conditions ~5.5 localizations/µm^2^ were obtained in regions outside cells suggesting that noise is about 16%. The entire time series was overlaid using a color-coded map or played as a video (Fig. 3a-c and video 3). At this temporal resolution, substantial changes in the localization of particles within the cell were observed, which did not allow following single localizations over time. Almost no movements were recorded for particles located outside cells, indicating that these changes do not result from drift (Fig. 3c, blue arrow, steady particle; red arrows moving particles). Although, individual shaker B channels could not be tracked over time a large-scale overview of the dynamic distribution of Shaker B channels at the PM could be obtained. To quantitatively estimate these large-scale changes, we plotted the distribution of molecule densities over time. Consistent with the SMLM data obtained in fixed cells the probability of a Shaker B channel to reside at low densities was the highest and did not change over time (up to ~100 particles/µm^2^), indicating that the overall density of Shaker B at the PM is relatively low and steady (Fig. 3d). The stable probability of Shaker B to reside at these densities throughout time further validates our assay conditions. Shaker B was additionally found to reside at higher densities (125-375 particles/µm^2^). At these high densities, however, the probability appeared to change between timepoints. These differences were irregular and fluctuated differently in different cells, indicating that they do not result from the acquisition itself but rather reflect a change in the probability of Shaker B to arrange at high-density units (Fig. 3d and S10). To confirm that these high densities indeed reflect specific organization of Shaker B at high densities, we performed a similar analysis on simulated data in which the same number of localizations, depicted at each timepoint, were randomly positioned inside the cell (Fig. S10b). The simulated data resulted in a steady probability to reside at low densities (up to ~100 particles/µm^2^) with zero probability to reside at the higher densities found in the data acquired in cells. Therefore, in cells, a small population of Shaker B appears to reside at high densities. Notably, while the low-density population was uniform and steady the high-density regions are heterogeneous and dynamic and may form and collapse within seconds. Shaker B is known to assemble into clusters at the PM upon binding to the scaffold protein PSD-95 ^29,30^. PSD-95 is specifically expressed in neuronal cells and therefore is not included in our heterologous COS7 expression system. The high-density regions observed here for Shaker B suggest that it can arrange in small unstable clusters at the PM in the absence of PSD-95.

**Fig. 3.**
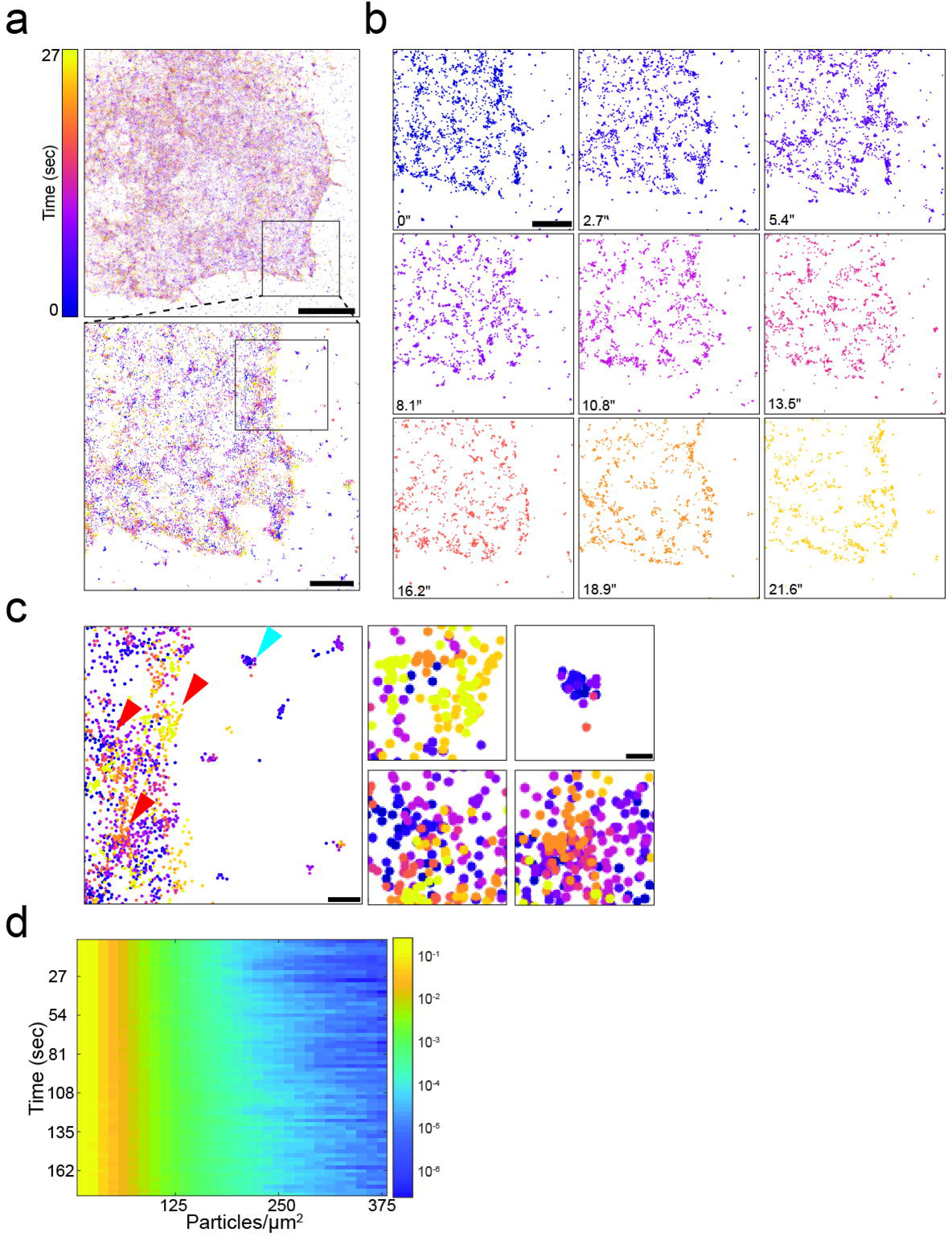
Bioorthogonal labeling enables super-resolution imaging of Shaker B in live cells. **(a)** A live-SMLM time stack image of a representative COS7 cell transfected with GCE system plasmid with Shaker B carrying BCNK via a peptide linker (Fig. S1a), labeled with 1.5 µM Tet-AF647. Timepoints are encoded by different colors. Each timepoint was reconstructed from 150 frames. A zoomed-in image of the region depicted in black square in top panel, is shown in bottom panel. Scale bars: top panel, 10 µm; bottom panel, 1 µm. **(b)** Time sequence of the cropped region shown in **a**. See also Video 3. Scale bar = 1 µm. **(c)** Left panel: zoomed-in image of the region depicted in black square in bottom panel in **a**. Right panels: zoomed-in images of regions located inside the cell (red arrows in left panel) and a region located outside of the cell (blue arrow in left panel). Scale bars: Left panel, 250nm; Right panels, 100 nm. **(d)** Three-dimensional density analysis of the cell shown in a-c; each row in the Y-axis represents a single timepoint, X-axis represents different particle densities and color encodes for the probability of a particle to reside at a given density, in logarithmic scale. Density profiles of additional cells and of simulated data, and a representative single timepoint plot are provided in figure S10.

### Sequential live-SMLM and SPT acquisitions in single cells via simultaneous two-color labeling of EFGR

In GCE-based bioorthogonal labeling of proteins, more than one type of Fl-dye can be applied at the labeling step, resulting in competitive labeling of the protein with different Fl-dyes ^6,41^. This can potentially be used for performing SPT and live-SMLM (that require Fl-dyes with different fluorescent properties) in the same cell. To this end, we expressed EGFR^128BCNK^ in COS7 cells and applied both Tet-Cy3 and Tet-AF647 at the labeling step. Cells were imaged live in SMLM mode, using AF647 channel, and then in SPT mode, using Cy3 channel. Using this protocol, both the overall distribution of molecules over time (live-SMLM) and the dynamics of EGFR molecules at the PM (SPT) were obtained from the same cell (Fig. 4a-d). Notably, such analysis allows correlating molecular densities and other spatial related information with diffusion rates at subregions of the cell, as demonstrated in figure 4e.

**Fig. 4.**
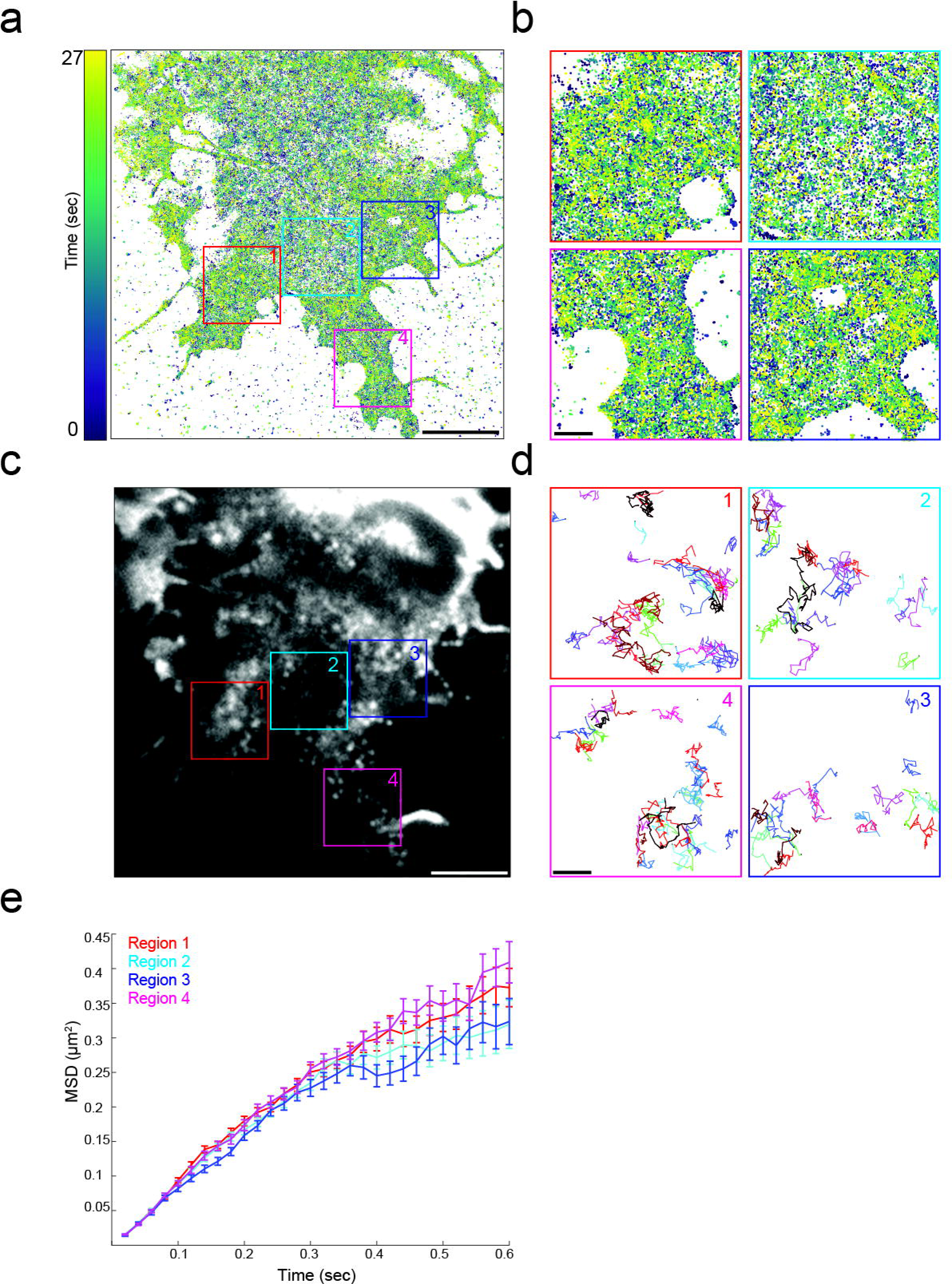
Performing single particle tracking and live-SMLM in the same cell by bioorthogonal labeling of EGFR with two Fl-dyes. COS7 cell transfected with GCE system plasmid and ERGR^128BCNK^ (Fig. S1a) and labeled with Tet-AF647 and Tet-Cy3 **(a)** A live-SMLM time stack image acquired in the 640 nm channel (Tet-AF647). Color represents single timepoints that were each reconstructed from 150 frames. **(b)** Zoomed-in images of four regions depicted in squares in **a**. **(c)** TIRF image of the same cell captured using 561 nm channel (Cy3), in SPT mode. **(d)** Tracks obtained from the zoomed in regions shown in **b**. Colors represent individual tracks. **(e)** MSD plots of tracks taken from each of the zoomed-in regions shown in **b** and **d**. Scale bars: a, c 5 µm; b, d 1 µm.

## DISCUSSION

To conclude, we showed that GCE-based bioorthogonal labeling of proteins with Fl-dyes could be used for live cell single molecule applications. SPT of EGFR and Shaker B carrying a ncAA and labeled with Tet-cy3, enabled measuring the diffusion of these proteins at different conditions. Moreover, live-SMLM were successfully performed and allowed recording the distribution of Shaker B at the PM over time with ~30 nm accuracy. Finally, by combining these two approaches in a single cell, the distribution of EGFR at the PM was documented at nanoscale resolution (live-SMLM) and its diffusion rates were measured with high accuracy (SPT). This opens up the opportunity to correlate protein diffusion with density using a minimal tag, that allows preserving the physiological properties of the proteins. Such information can shed light on the local spatiotemporal changes PM proteins undergo upon activation/inactivation conditions.

Diffusion rates ranging from 0.006 to 0.05 µm^2^/s were previously reported for EGFR using different labeling approaches and cell types, raising the possibility that the labeling method itself, the cell line or the analysis approach affect diffusion constants ^12,13^. In our study, stringent filtering was implemented to increase the reliability of our measurements, which can explain the faster diffusion rates measured in our study. Under these conditions, slower diffusion rates were measured for EGFR-Cy3 compared to EGFR-GFP (0.11 µm^2^/s and 0.15 µm^2^/s, respectively). Additionally, different brightness distributions were obtained for EGFR-Cy3 and EGFR-GFP in COS7 cells, suggesting that these molecules do not assemble at similar stoichiometries with the endogenous EGFR population. Given the ability of EGFR-Cy3 to assemble with endogenous EGFR at higher stoichiometries, the larger number of tracks and longer trajectories obtained for EGFR-Cy3, and the higher consistency with previously published data, we find the EGFR-Cy3 measurement better reflecting EGFR diffusion under physiological conditions. EGFR diffusion is known to be complex and to have more than a single diffusion component ^14^. By quantitatively characterizing EGFR diffusion we showed that EGFR diffusion is anomalous and confined. Notably, confined diffusion was seen in the presence or absence of the EGFR ligand, EGF, confirming previous reports of EGFR high-ordered organization prior to ligand activation ^12,14^. Consistent with previous findings, low doses of EGF did not affect EGFR diffusion on the PM while high doses slowed diffusion and increased confinement.

Measuring the diffusion of Shaker B channels using conventional N and C terminal tags is challenging due to the engagement of both tails of the protein in the activity of the channel ^31,32^. GCE-based labeling allowed overcoming these challenges by allowing labeling within the protein coding sequence. Additionally, the small size of the tag is expected to be exceptionally advantageous to Shaker B, as forming the functional channel pore relies on a complex protein-protein interaction network in the tetramer ^28^. The diffusion constants measured for Shaker B in our study are consistent with measurement performed for other voltage gated potassium channels (0.07 µm^2^/s vs 0.04 - 0.06 µm^2^/s) ^15,21^. In agreement with the tetrameric nature of the channels, diffusion rates obtained for Shaker B were slower than those measured for EGFR, which mostly resides in monomers and dimers ^14^. Moreover, Shaker B exhibited confined diffusion that was shown to be, at least partly, dependent on cortical actin. As experiments were performed in a heterologous system that does not express PSD-95, which presumably confines diffusion ^29,30^, these data indicates that Shaker B diffusion is confined even in the absence of a scaffold protein by additional cellular factors such as the actin cytoskeleton.

High density labeling is key for the success of super resolution microscopy techniques and specifically for SMLM. To apply these methodologies using GCE and bioorthogonal labeling, we performed a series of calibration and optimization steps. Under our optimized conditions, efficient labeling of EGFR and Shaker B at the PM was obtained (with higher labeling efficiencies obtained for EGFR) which allowed performing localization microscopy of these proteins in fixed and live cells. Background levels were relatively low for EGFR (6%) and marginal for Shaker B (20%). These differences in background levels most probably correspond to the different labeling efficiencies observed for these proteins. Under these conditions, live SMLM was performed for Shaker B at ~ 2.5 s temporal resolution. While background localizations (localizations measured outside the cell) appeared to be stationary, localizations inside the cell were very dynamic and exhibited density changes over time. Therefore, performing live-SMLM on bioorthogonally labeled PM proteins can be used for recording the dynamics of proteins at the PM with high accuracy. Notably, the temporal resolution used here was determined based on the number of localization obtained in a single frame and the maximal frame rate of the camera (55 fps). Temporal resolution may be further improved by increasing labeling densities or by using higher frame rate cameras.

SPT and SMLM were previously performed on bioorthogonally labeled glycans by hijacking the metabolic pathway of glycan synthesis and introducing artificial monosaccharides that carry a bioorthogonal handle ^42,43^. As numerous PM proteins are known to be glycosylated, these studies provide a unique view on the overall organization of proteins on the PM. In these studies, the diffusion coefficient values were between 0.06 to 0.13 µm^2^/s, which is in the range of the values we obtained for EGFR and Shaker B, confirming our SPT assay and analysis ^42^. Homogenous distribution of glycans at the PM was observed in neuroblastoma and U2OS cells, using SMLM ^43^. Together with the overall homogenous distribution we observed using SMLM for EGFR and Shaker B on the PM, these data support the notion that proteins are uniformly distributed on the PM and are not organized in stable domains with a defined size. However, this does not rule out the existence of small, dynamic clusters as inferred from our SPT and live-SMLM experiments.

The high labeling efficiencies obtained for EGFR^128BCNK^ enabled SPT and live-SMLM to be recorded in the same cell. Because our microscope setup includes one camera, these datasets were recorded sequentially. Using a dual cam datasets may be recorded in parallel, providing the opportunity to document both the distribution and the dynamics of PM proteins in real time.

## MATERIALS AND METHODS

### Cell culture

COS7 cells were grown in Dulbecco’s Modified Eagle Medium supplemented with 10% fetal bovine serum (FBS), 2 mM glutamine, 10,000 U/ml penicillin, and 10 mg/ml streptomycin.

### Genetic code expansion and bioorthogonal labeling in mammalian cells

For live cell experiments, cells were plated 24 h before transfection at 15% confluency in a four-well µ-Dish glass-bottom chamber slide (Ibidi, Martinsried, Germany). Transfection with the plasmids described in figure S1a, was carried out using Lipofectamine 2000 (Life Technologies, Carlsbad, CA) according to the manufacturer’s guidelines. Genetic code expansion and biorthogonal labeling was carried out as described in Schvartz et al. ^11^. In short, cells were incubated for 48 h in the presence of BCNK (0.5 mM; Synaffix, Oss, Netherlands) and washed with fresh medium at 37°C. Labeling was then carried out in the dark (30 min 37°C) with 1.5 µM of: Tet-Cy3, Tet-Cy5, Tet-ATTO532 (Jena Bioscience GmbH, Jena, Germany) or AF647 (Thermo Fisher Scientific, Waltham, MA), that was coupled with the tetrazine as previously described ^9^. For live cell TIRF imaging, all cells were kept at 4°C to avoid internalization Finally, cells were washed and imaged as described below.

### Live cell imaging

Single Z-slices were collected at 37°C using a fully incubated confocal spinning-disk microscope (Marianas; Intelligent Imaging, Denver, CO) with a 63X N.A 1.4 oil immersion objective and were recorded on an electron-multiplying charge-coupled device camera (pixel size, 0.079 µm; Evolve; Photometrics, Tucson, AZ). Image processing and analysis were done using SlideBook version 6 (Intelligent Imaging).

### Single particle imaging and analysis

Images were collected on the Zeiss Elyra inverted wide field fluorescence microscope using a Zeiss 100X N.A 1.46 TIRF oil immersion objective. Excitation was achieved using solid-state lasers (488 nm for GFP or 561 nm for Tet-Cy3 and mCherry). Laser intensities and exposure times were optimized in each labeling strategy for maximal SNR and minimal photobleaching. Images were recorded on an electron-multiplying charge-coupled device camera (iXon DU897, Andor, Belfast, Northern Ireland). For each SPT experiment at least 500 images and up to 10,000 were recorded with a frame rate of 50 fps. For Cy3 labeled samples (EGFR-Cy3, Shaker B-Cy3) a short (~10 s) bleaching step with laser intensities of ~20% was applied prior to acquisition to reduce particle density and enable single foci detection. For EGF stimulation cells were maintained in FBS free media for 2 hours prior to imaging followed by the addition of 10 nM, 100 nM recombinant human EGF (Sigma-Aldrich, St. Luis, MS). For LatA treatments 1 µM LatA was added to cells and incubated for 10 min at 37°C prior to imaging. SNRs were determined for individual spots by measuring the intensity profile along a line that crosses a few spots. Peak intensities were regarded as signal while the average local minima between the peaks was regarded as noise. Three time-points were chosen in each acquisition and 4-5 particles were measured in each one. This was repeated for 10 cells from each sample (EGFR-Cy3 or EGFR-GFP). Brightness analysis was performed by plotting the distribution of intensity values obtained for each particle, as measured in the second frame of the time acquisition. Plots were fitted with a combination of Gaussian distributions. Best fits are shown in supplementary figures S2, S6a.

### Single particle tracking

Movies of cells, recorded on different days, were quantified for each condition. Single particles were detected and tracked using a custom MATLAB (MathWorks, Natick, Massachusetts) code. The ensemble average mean squared displacement of traced particles was defined as the average over all particles of the squared displacement of each particle at time *τ* from its position at *τ* = 0:

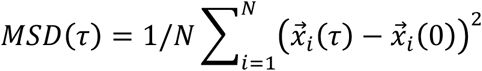

where 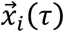 is the position of particle *i* at time *τ*, and *N* is the total number of tracked particles.

Background filtering was performed by measuring diffusion of nonspecific Fl-dye molecules attached to the glass bottom and including and excluding tracks with similar diffusion rates, as detailed in figure S3. In addition to background filtering, trajectories were filtered based on length; only trajectories longer than 20 frames were considered. To avoid bias due to difference in trajectory lengths, MSD values for filtering purposes were calculated only for the first 20 frames of each trajectory.

The distribution of the diffusion coefficients was calculated using the time averaged MSD of each trajectory:

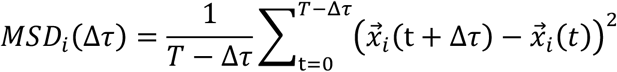

where i indicates a particle index and T is the total number of frames the particle was tracked.

The diffusion coefficient for each particle is then derived from the slope of the *MSD*_*i*_(Δ*τ*) for Δ*τ* ≤ 0.1*sec*, which is the linear domain of the MSD, according to 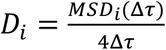. We have obtained the slope by linear least square fit and considered only results that had less than 0.01 standard regression error. We note that the ensemble and time average MSDs do not coincide, which is typical for protein diffusion in the PM ^22^. The turning angle is defined as the angle between the direction of a displacement 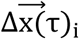 and a following displacement 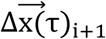. Probability distribution of turning angles was calculated as previously described ^23^.

### Fluorescence recovery after photobleaching (FRAP)

Time series were collected on a Zeiss LSM880 inverted laser scanning confocal microscope using a Zeiss 63X N.A 1.4 oil immersion objective. COS7 cells expressing EGFR-Cy3 were imaged for 5 timepoints (pre-bleach), followed by bleaching a specified ROI along the membrane contour. Cells were then imaged (post bleach) until fluorescence intensity reached a plateau. Intensity levels were subjected to background subtraction and photobleaching correction. Experimental results were fitted to a single exponential decay curve using GraphPad (GraphPad Software, CA) from the fit the half-life of recovery (*t*_1/2_) was obtained. Using the half-life value, the diffusion coefficient was calculated as follows: 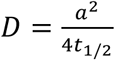 where a represents the width of the area bleached as described in ^44^.

### Single Molecule Localization Microscopy (SMLM)

Images were collected on the Zeiss Elyra inverted wide field fluorescence microscope using a Zeiss 100X N.A 1.46 TIRF oil immersion objective as described in live cell single molecule section. All experiments were performed using uHP laser focusing mode available in the Zeiss Elyra system. Fixed samples were imaged using the following buffer: 50 mM TRIS, 10 mM NaCl to pH=8.0 supplemented with 10% glucose (%W/V), GLOXY oxygen scavenging system (170 U/mL Glucose oxidase, 1,400 U/mL Catalase) and 10 mM mercaptoethylamine (MEA) ^37^. Live samples were imaged in OptiMEM™ (Gibco™; Dublin; Ireland) supplemented with 2 mM glutamine, 5% glucose (%W/V), GLOXY oxygen scavenging system (170U/mL Glucose oxidase, 1,400U/mL Catalase) and 1% β-mercaptoethanol (%V/V) (BME). For fixed cells, 100% laser power was used, and 405 nm irradiation was applied as needed. For live cells, 70% laser power was used, and 405 nm irradiation was applied in linearly increasing amounts up to a maximum of 5%. For both fixed and live cell SMLM, 10,000 images were collected in each experiment with a frame-rate of 55 fps using irradiation intensities of ~1-3 kW/cm^2^.

### SMLM image analysis

Single particles were localized, fitted and reconstructed using ThunderSTORM ^45^. Fixed SMLM localizations were filtered by number of photons and localization uncertainties. Localization uncertainty was calculated in ThunderSTORM as described in ^45^. In short, uncertainty values were calculated for each localization spot based on the standard deviation of the Gaussian function fitted to each spot, the back-projects pixel size, and the number of photons.

Images were drift corrected using cross-correlation prior to reconstruction. In live SMLM, localizations were filtered, but not drift corrected. Localizations obtained from 150 sequential frames were compiled into a single timepoint, using a custom written Python code (see optimization in figure S9). Each frame was acquired at 18 ms. Density analysis (Fig. 4d and S10) was performed for each timepoint using a custom written Python code, by applying a circular rolling window with a 50 nm radius. Number of particles was normalized for each time point. All codes are freely available at https://gitlab.com/sorkin.raya/storm-analysis. Randomized particle distribution simulation: in each stack of 150 frames, the total number of particles was counted. Then, the same number of particles was randomly positioned within the cell, with arbitrarily generated x,y coordinates, thus creating a randomized distribution of particles. A mask to identify the cell location was created based on the density of particles summed over the whole movie series. Density analysis of the particle distribution was then performed as described above.

### Live SMLM-SPT imaging

COS7 cells were transfected as described above and bioorthogonally labeled with both Tet-Cy3 and Tet-AF647 (150 nM and 1.5 µM respectively). Cells were first imaged in live-SMLM as described above (5,000 frames) using a 640 nm channel. Then, live-SMLM buffer was substituted with growth media and the same cell was imaged using the SPT protocol described above. Videos were then processed for both live-SMLM and SPT as detailed above.

## AUTHOR CONTRIBUTIONS

A.K and N.E designed and were involved in the analysis of all experiments. A.K performed all experiments. A.K and A.A helped writing the manuscript. D.N and E.A provided technical help and knowledge in genetic code expansion. K.D and E.A synthesized Tet-AF647. R.S and Y.R did all SPT analysis and measurements. R.S and A.K generated the figures and sup figs. G.B and R.S performed density analysis in live-SMLM. O.Y contributed Shaker B plasmid and helped designing Shaker B mutants. N.E conceptualized the work and wrote the manuscript.

## FUNDING SOURCES

The project leading to this application has received funding from the European Research Council (ERC) under the European Union’s Horizon 2020 research (N.E, startup grant). Y.R is supported by the ISF (Israel Science Foundation) grant 988/17. R.S acknowledges support through HFSP postdoctoral fellowship LT000419/2015.

## ACKNOWLEDGMENTS

We thank Levi Gheber for help with FRAP analysis and Ronit Pinkas-Kramarski (TAU, Israel) for contributing the EGFR plasmid. We also thank Dr. Uzi Hadad from the Ilse Katz Institute for Nanoscale Science and Technology Shared Resource Facility for technical help with Zeiss LSM880 Airyscan confocal system. Finally, we thank members of the Elia laboratory for fruitful discussions and critical reading of the manuscript.

