## Supplementary Figures for "Live cell single molecule tracking and localization microscopy of bioorthogonally labeled plasma membrane proteins"

Supplementary figure 1

a

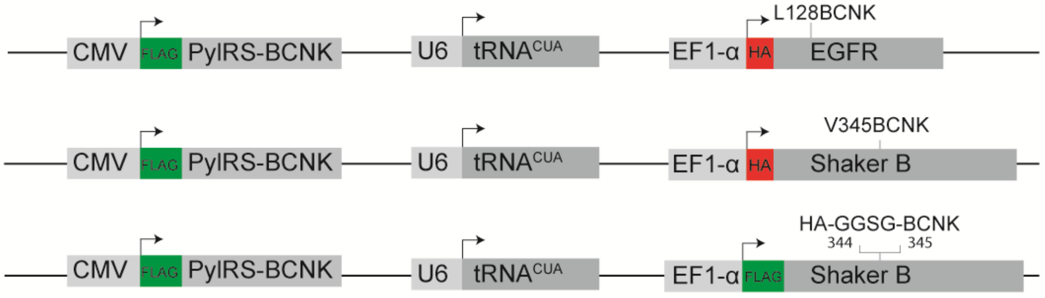


b

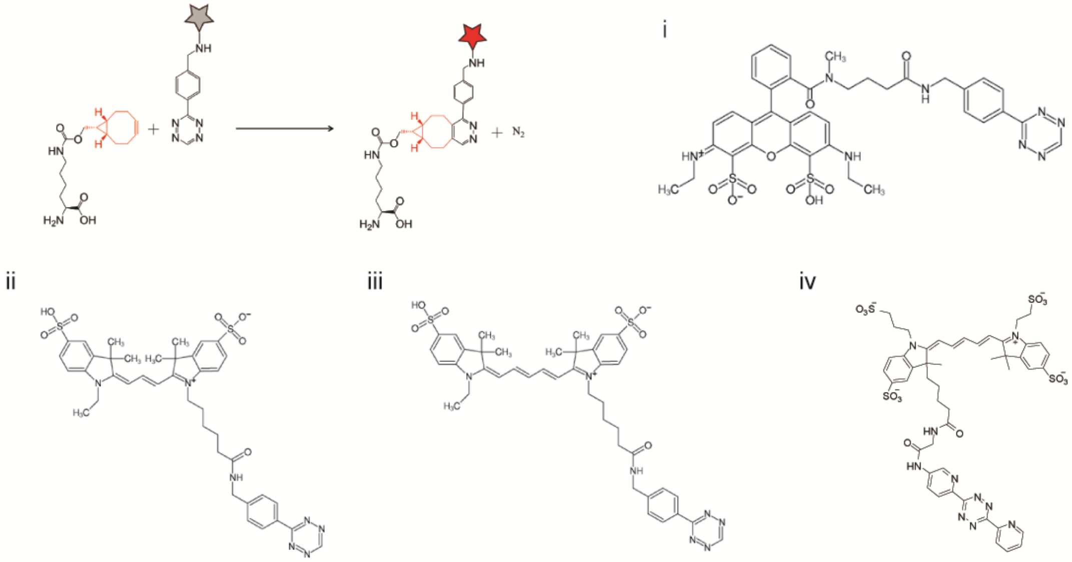

c

| Fl-dye | Intensity (a.u) | Photobleaching (%intensity/sec) | Track lenght (frmaes) | Number of tracks |
|---|---|---|---|---|
| Tet-Cy3 | 44.6±6.6 | 0.162±0.008 | 23 (11, 115) | 1,951±233 |
| Tet-Cy5 | 20.8±4.8 | 0.164±0.009 | 22 (11, 99) | 907±328 |
| Tet-AF647 | 18.3±2.8 | 0.105±0.009 | 21 (11, 88) | 554±26 |
| Tet-ATTO532 | 17.2±2.5 | 0.122±0.014 | 17(10, 49) | 1,329±134 |

**Supplementary Fig. S1. Single vector GCE expression constructs and *in-vitro* evaluation of Tet-Fl-dyes for SPT.** GCE plasmid constructs used in this work. All constructs contain a single copy of an evolved Pyrrolysine tRNA synthetase (PylRS) that aminoacylates BCNK, expressed under a CMV promoter and a single orthogonal amber suppressor $tRNA^{CUA}$ expressed under a U6 promoter. All proteins of interest are expressed under an EF1- α promotor. Top panel: EGFR with an in-frame stop codon at position 128. Middle panel: Shaker

B channel with an in-frame stop codon at position 345. Bottom panel: Shaker B channel with a short linker inserted after position 344, the linker contains an HA tag followed by a short Gly-Gly-Ser-Gly linker and finally a TAG codon encoding for BCNK. **(b)** Inverse-electron demand Diels-Alder reaction between Bicyclo[6.1.0]nonyne-L-lysine, a modified lysine ncAA carrying a strained Alkyne and a Fl-dye conjugated to a tetrazine. Resulting in the formation of a covalent bond between the ncAA and the Fl-dye with nitrogen as the only byproduct (top left panel). Tet-Fl-dye conjugates used in this work: Tet-ATTO532 (i), Tet-Cy3 (ii), Tet-Cy5 (iii) and Tet-AF647 (iv) **(c)** Comparing several photophysical properties of Fl-dye tetrazine conjugates. Fl-dyes were plated on a glass-bottom coverslip in the presence of 0.5 mM BCNK and imaged in TIRF mode using the same conditions described for SPT acquisition in methods section. Parameters measured included: mean intensity (a.u, ±sd), photobleaching (%intensity/s, ±sd), median track length (number of frames with 95% confidence interval) and the mean number of tracks per acquisition (±sd) (n=5 independent experiments).

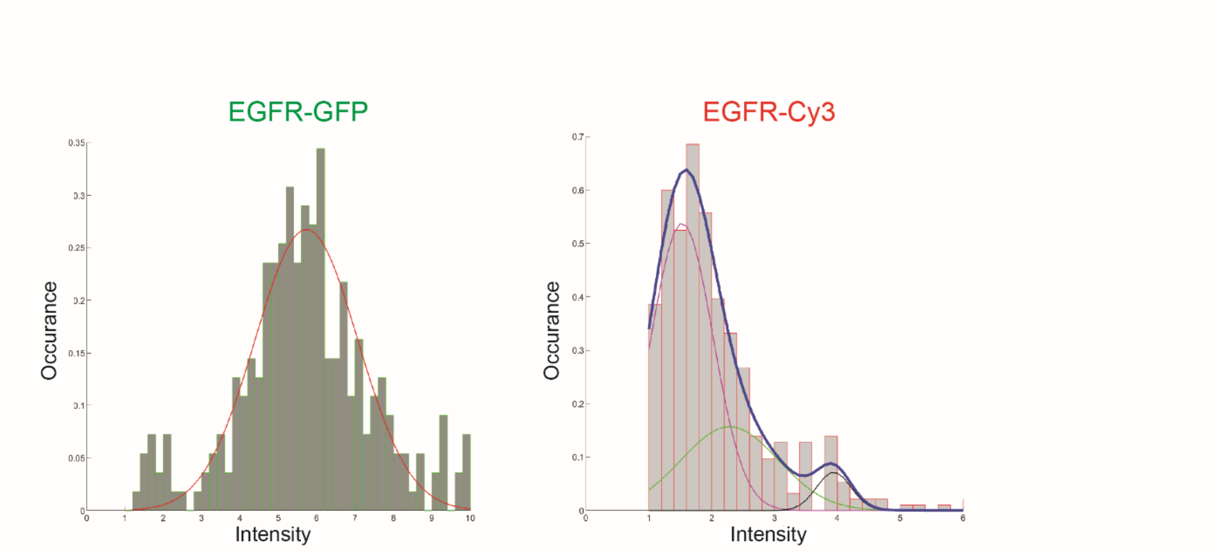


**Supplementary Fig S2.** Intensity value distributions of the second frame of all SPT experiments performed with either EGFR-GFP (left panel) or EGFR-Cy3 (right panel), fits correspond to single or double gaussian fits. ($R^2_{EGFR\text{-}GFP}$= 0.98, $R^2_{EGFR\text{-}Cy3}$= 0.99)

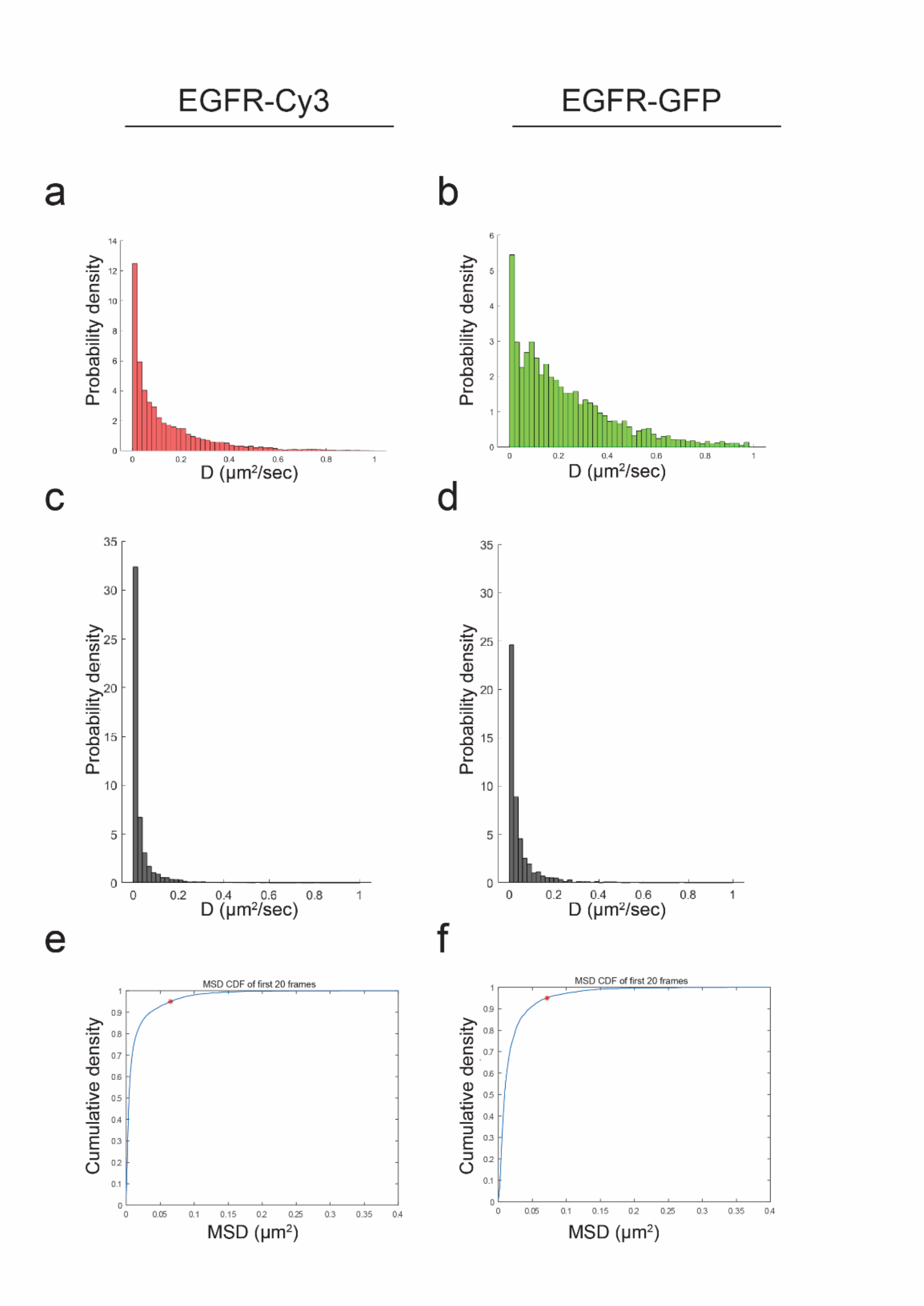


**Supplementary Fig S3. Background filtering in SPT analysis of EGFR. (a, b)** Diffusion coefficients [D] distributions of EGFR-Cy3 and EGFR-GFP respectively without filtering. **(c, d)** Diffusion coefficients [D] distributions of Tet-Cy3 and GFP molecules respectively that were located outside cells. **(e, f)** Cumulative distribution of mean square displacement (MSD) of background values presented in S2C and S2D, respectively. Red stars indicate the top 5 percent of MSD chosen as the threshold for filtering: 0.065 $\mu m^2$for Cy3 (n=5 independent experiments) and 0.07 $\mu m^2$ for GFP (n=5 independent experiments).

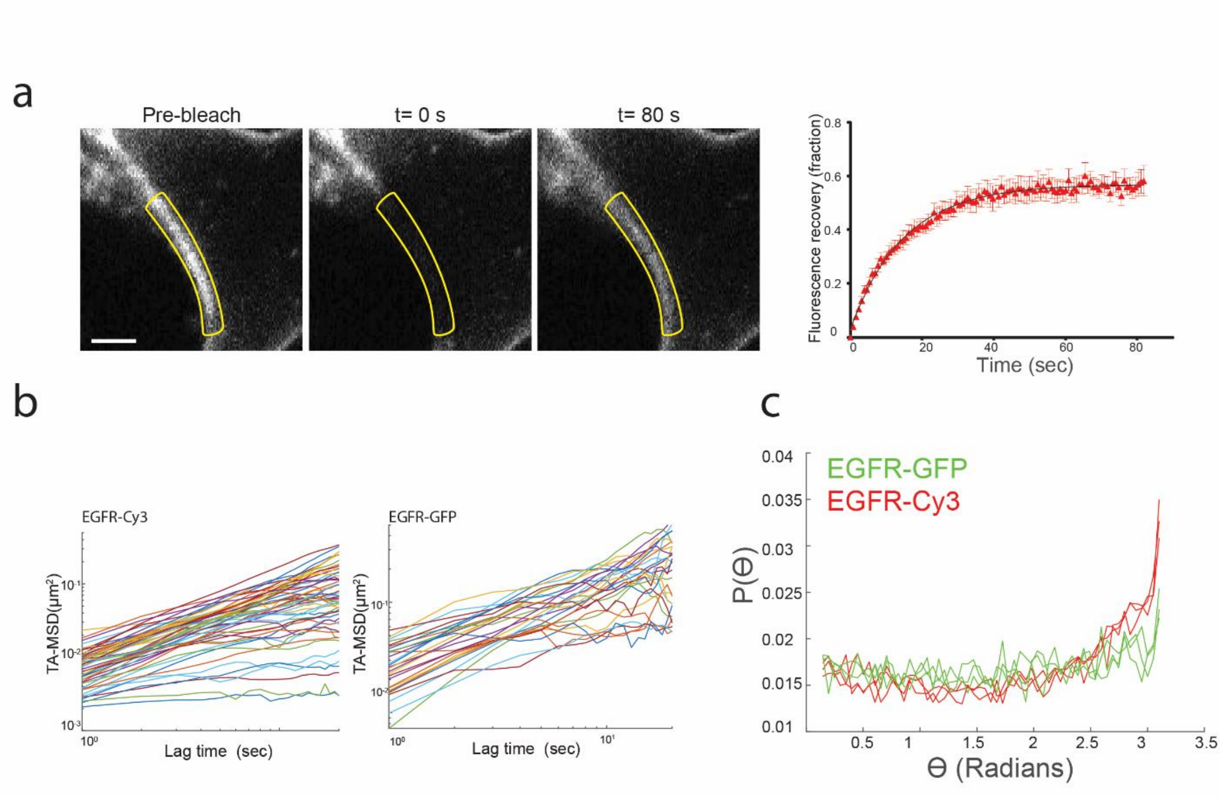


**Supplementary Fig S4 (a)** COS7 cell expressing EGFR$^{L128BCNK}$ and labeled with Tet-Cy3. First panel: a pre-bleached image. Second panel: one frame after photobleaching. Third panel: image acquired 80 seconds post photobleaching, showing recovery. Photobleached ROI is marked in yellow, Scale bar = 2 μm. Forth panel: ensemble of fluorescence recovery plots used to extract the diffusion coefficient as described in methods sectiond. Error bars represent standard deviations (D=0.05 μm$^2$/sec n=5, $t_{1/2}$ = 11.26 $\pm$ 2.96 s). **(b)** Time-averaged MSD (TA-MSD) as a function of lag time for individual trajectories. The variability intersection point of the individual particle MSDs points to non-ergodicity and heterogeneity of the membrane. **(c)** Turning angle distributions constructed for lag times of 1,2 and 5 frames.

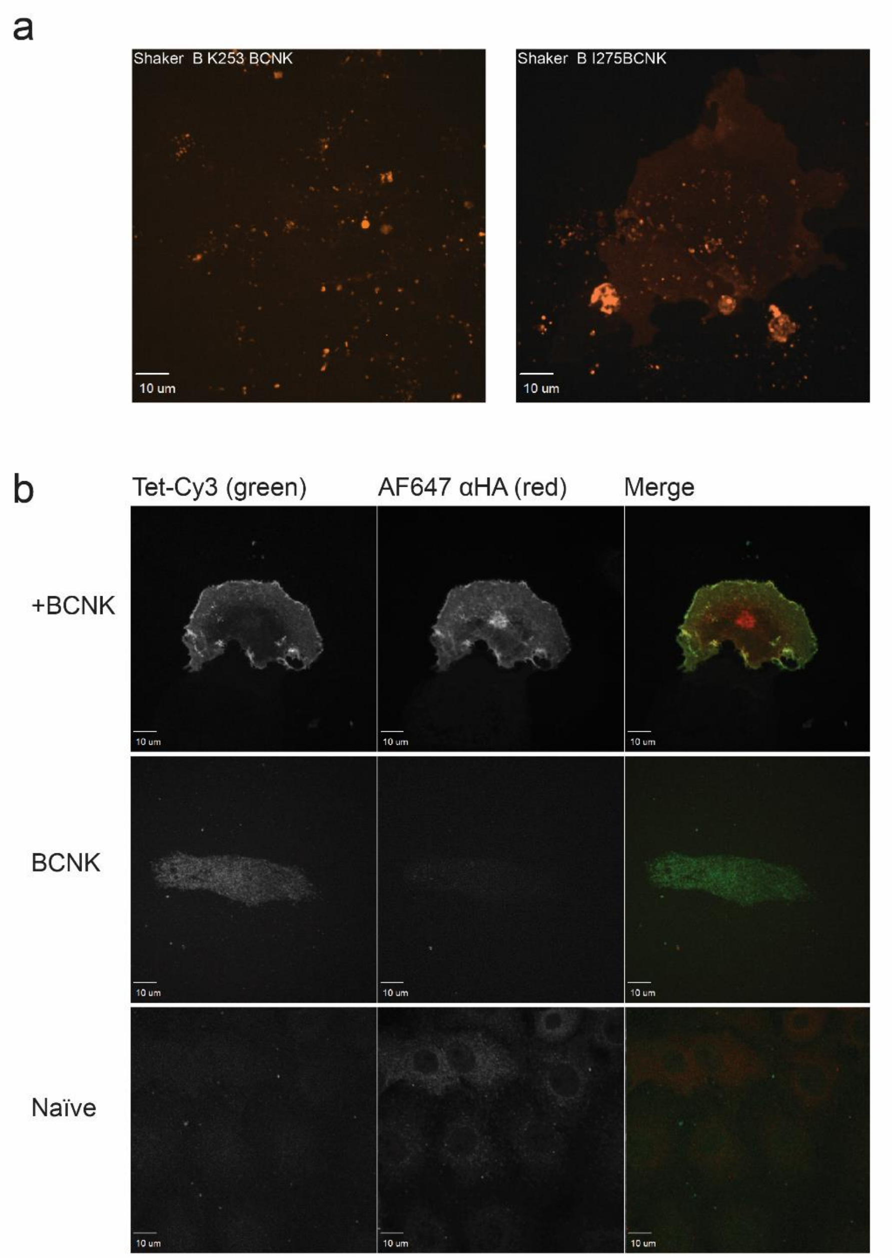


**Supplementary Fig S5. Evaluating different positions for the bioorthogonal labeling of Shaker B. (a)** Fluorescence spinning disk microscope images of cells expressing the GCE system and Shaker B with an in-frame stop codon at either position 253 (left panel) or 275 (right panel) bioorthogonaly labeled with 1.5 μM Tet-Cy3. Little to no labeling was observed using Shaker $B^{K253BCNK}$ and Shaker $B^{I275BCNK}$ **(b)** Fluorescence spinning disk microscope images of naïve cells or cells expressing the GCE system and Shaker B with an in-frame stop codon at the optimized position (345) in the presence (+BCNK) or absence (-BCNK) of BCNK. Cells were labeled with 1.5 μM Tet-Cy3, fixed and immunolabeled with rabbit α-HA followed by a secondary antibody conjugated to AF647. Detection was made at 561nm wavelength for Tet-Cy3 (left) or 647 nm wavelength for HA-AF647 (middle). A merged channel demonstrating co-localization is shown to the right. This demonstrates robust and specific bioorthogonal labeling of Shaker $B^{V345BCNK}$.

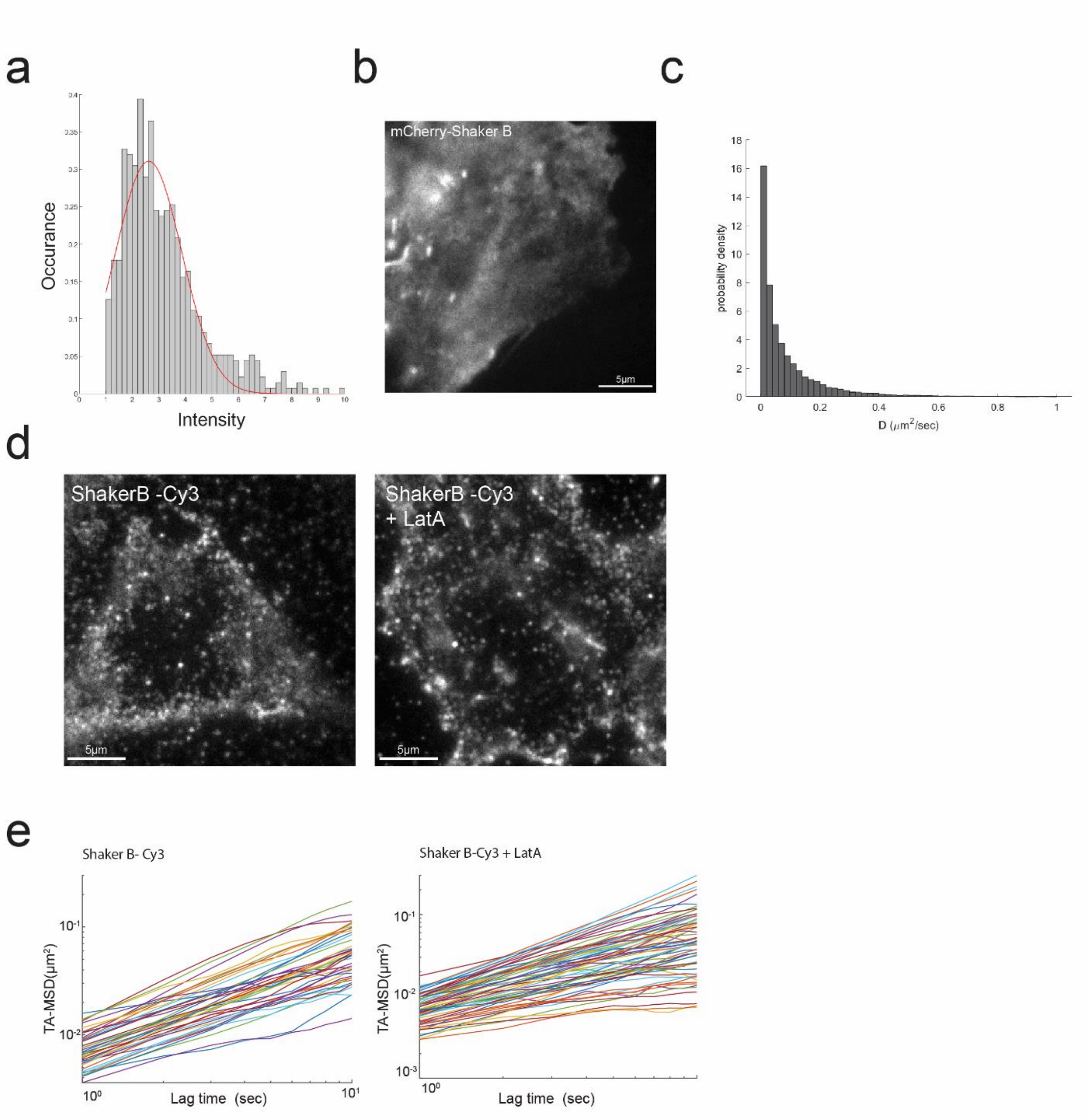


**Supplementary Fig S6. SPT analysis of bioorthogonally labeled ShakerB. (a)** Intensity value distribution at the second frame of all SPT experiments performed with Shaker B-Cy3, fit corresponds to a single gaussian ($R^2 = 0.9884$). **(b)** A representative TIRF image of a COS7 cell expressing mCherry-ShakerB. **(c)** Unfiltered distribution of diffusion coefficients [D] of Shaker B-Cy3. **(d)** Representative TIRF images of COS7 cells expressing ShakerB$^{linkerBCNK}$ and labeled with Tet-Cy3 without (left panel) or with (right panel) 1μM LatA treatment. **(e)** Time-averaged MSD (TA-MSD) as a function of lag time for individual trajectories. The variability intersection point of the individual particle MSDs points to non-ergodicity and heterogeneity of the membrane.

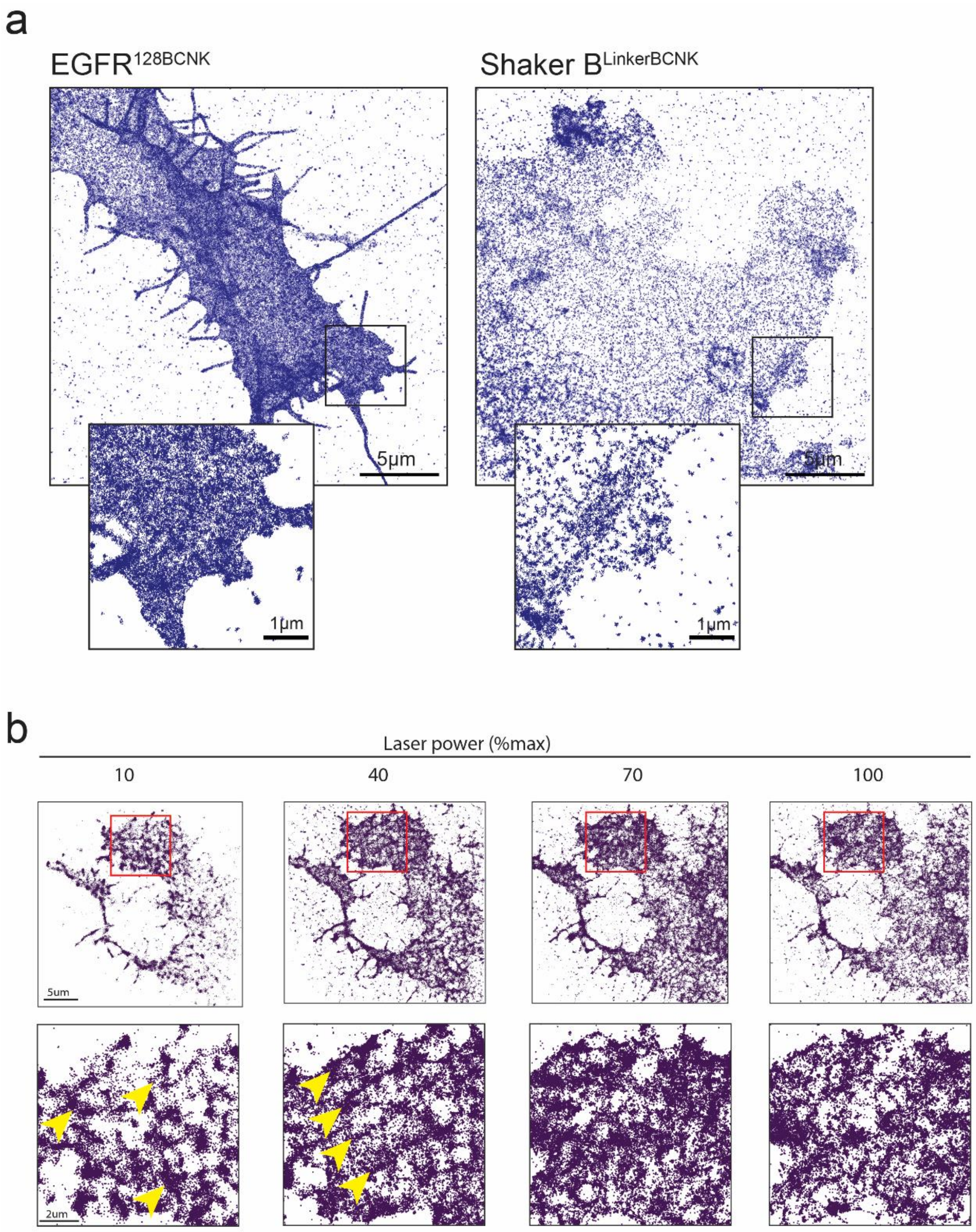


**Supplementary Fig S7. SMLM in fixed cells. (a)** Fixed COS7 cells expressing either EGFR$^{L128BCNK}$ (left panel) or ShakerB$^{linkerBCNK}$ (right panel) were labeled with 1.5μM Tet-AF647 and subjected to SMLM imaging. Insets are zoomed-in images of the region labeled in black-square in each image. (n for EGFR$^{L128BCNK}$=7, n for ShakerB$^{linkerBCNK}$ =3). **(b)** Fixed COS7 cell expressing EGFR$^{L128BCNK}$ and labeled with 1.5 μM Tet-AF647 were subjected to SMLM imaging using different excitation laser powers (percent of maximal laser power). A total of 2000 frames were acquired for each SMLM image. Bottom panels are zoomed-in images of the areas labeled in red squares in upper panels. Yellow arrows mark typical artifacts caused by insufficient laser intensities.

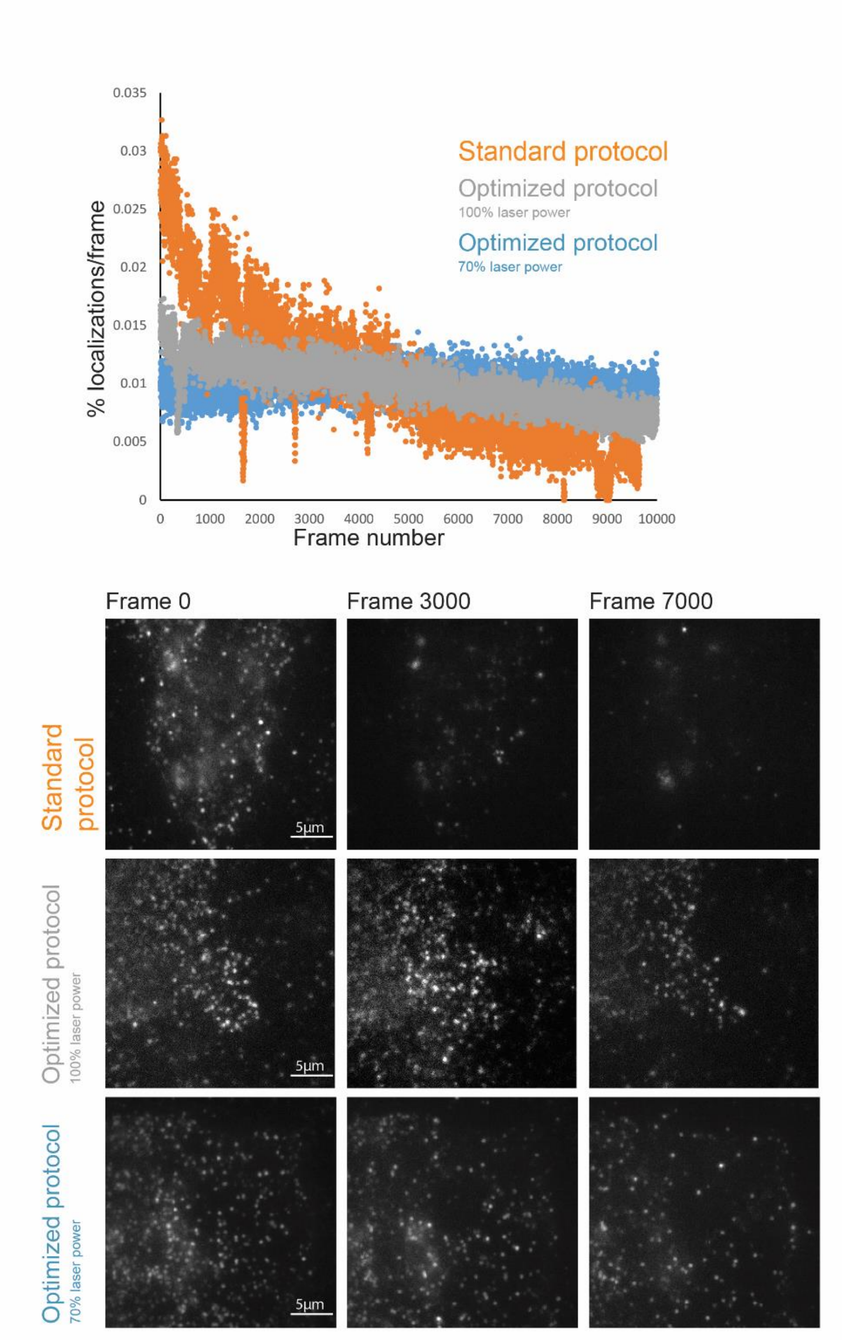


**Supplementary Fig S8. Optimized acquisition protocol for live SMLM.** Scatter plots of the percentage of localizations per-frame obtained in a live cell expressing ShakerB$^{linkerBCNK}$ -Cy3 as a function of frame number for a standard SMLM protocol (orange dots), an optimized protocol using maximal laser power (grey dots) and an optimized protocol using 70% laser power (blue dots). Standard SMLM protocol consists of standard imaging buffer and acquisition setup described for fixed cells in methods section. Optimized protocol consists of optimized live-SMLM buffer and acquisition parameters described in methods section for live –SMLM. Bottom panel are representative TIRF images taken from frames 0, 3000 and 7000 of the cells plotted in top panel.

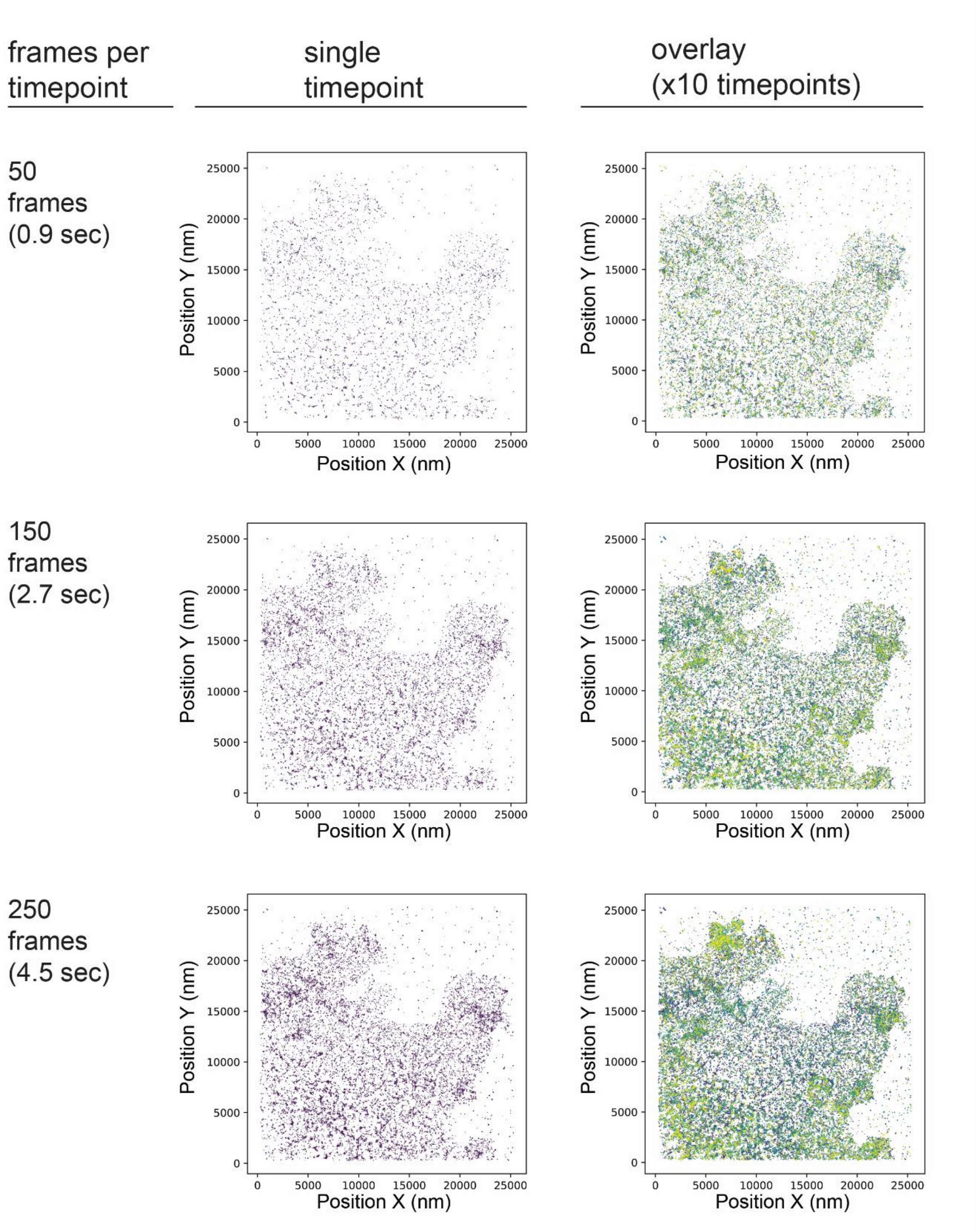


**Supplementary Fig S9. Estimating temporal resolution for live-SMLM.** Fixed COS7 cells expressing the GCE system plasmid with ShakerB$^{linkerBCNK}$ were labeled with 1.5µM Tet-AF-647, fixed and subjected to SMLM imaging. Temporal resolution was estimated by overlaying localizations obtained from 50 frames (top panel), 150 frames (middle panel) and 250 frames (bottom panel), resulting in a temporal resolution of 0.9 sec, 2.7 sec and 4.5 sec, respectively. Shown from left to right: number of frames overlaid, an image of the first timepoint, an overlay of 10 time-points (color-coded for time).

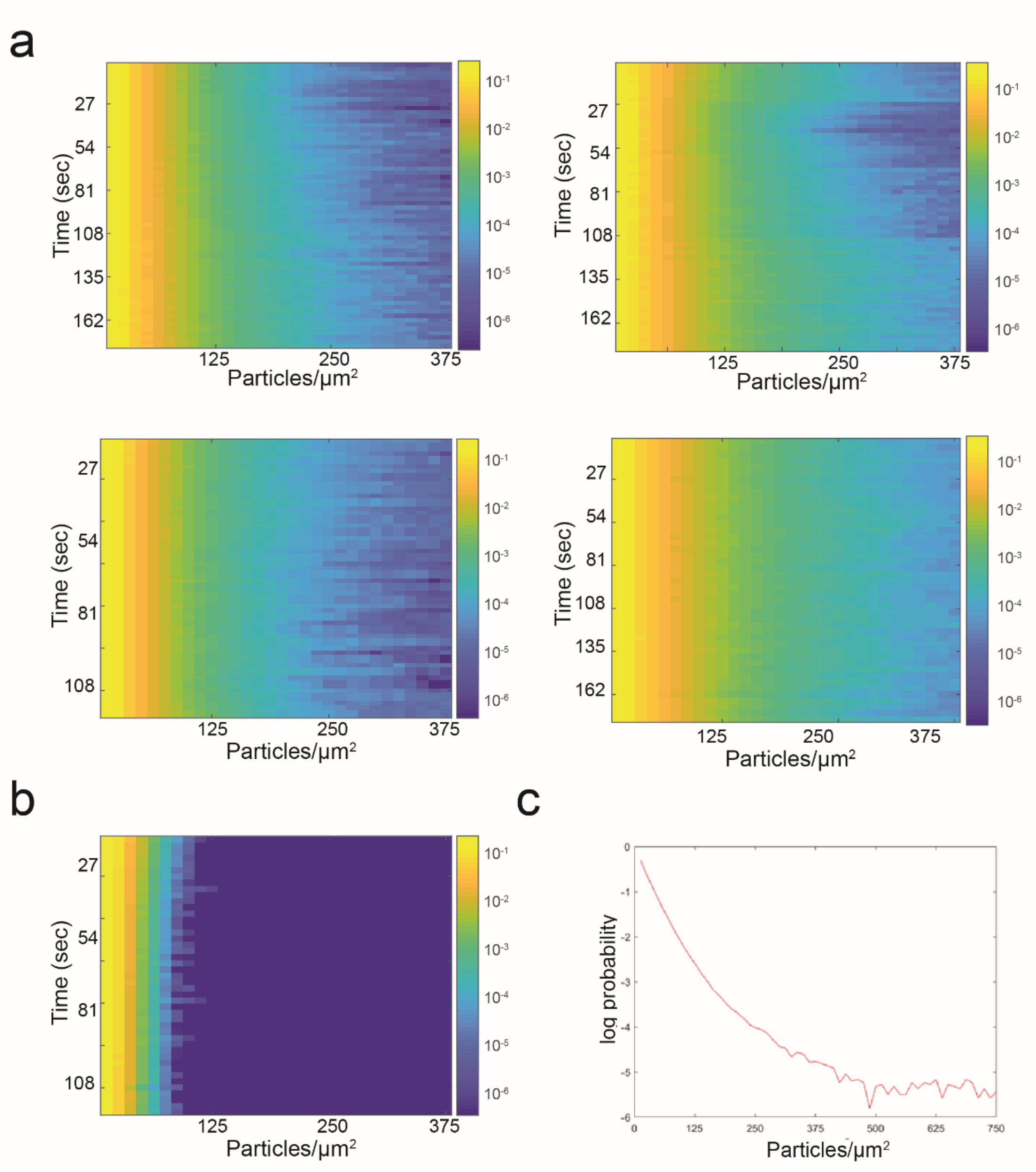


**Supplementary Fig S10. Density-probability plots obtained from live-SMLM experiments.** **(a)** Probability plot of particle density per timepoint. Each point on the Y-axis represents a single 150 frame reconstruction, X-axis are different particle densities and the color encodes for the log probability of a particle to be at a specific density. From each such timepoint, the probability distribution function is calculated and presented on a logarithmic scale. Probability density for each time point (150 frames) was calculated as a function of particle density. Each plot was generated from a different cell and represents a different live-SMLM experiment. Total number of localizations obtained in the first timepoint were: top left, 42403; top right, 26967; bottom left, 28034, bottom right: 25600. **(b)** Probability plot of particle density per timepoint obtained for simulated data. Simulation was performed by taking the total number of localizations obtained for the cell presented in figure 3d at each timepoint and placing them randomly inside the cell. **(c)** Log probability curve obtained for a single timepoint.
